# Contribution of UbrA, a ubiquitin ligase essential for Arg/N-degron pathway, to peptidase gene expression in *Aspergillus oryzae*

**DOI:** 10.1101/2025.04.16.649092

**Authors:** Waka Muromachi, Mao Ohba, Yasuaki Kawarasaki, Youhei Yamagata, Mizuki Tanaka

**Author notes:** Address correspondence to Mizuki Tanaka,.

## Abstract

The degradation of intracellular proteins by N-degron pathways depends on their N-terminal amino acids. In budding yeast, the Arg/N-degron pathway controls the expression of di/tri-peptide transporter gene by degrading transcriptional repressor. However, there is no detailed information on the N-degron pathway in filamentous fungi, and its role in regulating microbial nitrogen metabolism is unclear. Here, we demonstrated that the E3 ubiquitin ligase, UbrA, which is required for the Arg/N-degron pathway, regulates peptidase gene expression in the filamentous fungus *Aspergillus oryzae*. Using ubiquitin-fused green fluorescent protein as a reporter, we showed that the Arg/N-degron pathway in *A. oryzae* is similar to that in budding yeast. Disruption of *ubrA* significantly reduced activities of acidic endopeptidase and carboxypeptidase in submerged culture using soy protein as the nitrogen source. In addition, *ubrA* disruption dramatically reduced the mRNA expression of the major endopeptidase and carboxypeptidase genes but increased alkaline peptidase production. Moreover, *ubrA* disruption reduced the expression levels of di peptidyl- and tripeptidyl- peptidase genes and di/tri-peptide transporter genes. This regulation was independent of PrtR, the transcription factor of major extracellular peptidase genes. Our data showed that UbrA is involved in the expression of various peptidase genes in concert with di/tri-peptide transporter genes.

**IMPORTANCE:** Peptidases produced by *Aspergillus oryzae* are important in the production of Japanese fermented foods and are used as industrial enzymes for various food-processing and pharmaceutical applications. The expression of di/tri-peptide transporter gene in budding yeast is controlled by a positive feedback mechanism through the dipeptide-mediated activation of the E3 ubiquitin ligase, Ubr1, which is essential for the Arg/N-degron pathway, which determines the lifetime of intracellular proteins. In this study, we demonstrated that *A. oryzae* UbrA (an ortholog of yeast Ubr1) regulates peptidase gene expression in addition to di/tri-peptide transporter genes. Disruption of *ubrA* decreases the expression of major acidic peptidase genes and increases the expression of alkaline peptidase gene. In addition, the expression levels of di/tri-peptidyl peptidase genes and di/tri-peptide transporter genes were reduced by *ubrA* disruption. These results suggest that UbrA regulates the expression of various peptidase genes to facilitate positive feedback of di/tri-peptide transporter genes.

## INTRODUCTION

Most proteins undergo post-translational processing, such as methionine removal by methionine aminopeptidase (MetAP), cleavage by endopeptidase, and deamidation or arginylation of N-terminal amino acids. The lifetimes of intracellular proteins depend on their N-terminal amino acid residues. In eukaryotic cells, the recognition of N-terminal amino acids by ubiquitin ligases determines the stability of intracellular proteins. This system is called the N-degron pathway (or N-end rule) (1, 2). In *Saccharomyces cerevisiae*, the E3 ubiquitin ligase, Ubr1, directly recognizes the N-terminal amino acid residues at two distinct substrate-binding sites, type-1 and type-2 (3, 4). The type-1 site recognizes the basic amino acids arginine, lysine, and histidine. The type-2 site recognizes the bulky hydrophobic amino acids leucine, phenylalanine, tryptophan, tyrosine, isoleucine, and unacetylated methionine. Proteins with N-terminal asparagine, glutamine, aspartate, and glutamate are also recognized by Ubr1 because the amido group of N-terminal asparagine and glutamine residues is removed by amidase, and arginine is conjugated to N-terminal aspartate and glutamate by arginyl-tRNA-protein transferase (1, 2). The proteins recognized by Ubr1 are ubiquitinated and rapidly degraded by the proteasome. This Ubr1-mediated degradation pathway is called the Arg/N-degron pathway. When the N-terminal methionine itself or N-terminal alanine, serine, threonine, valine, cysteine, and glycine exposed after methionine removal by MetAP are acetylated by the ribosome-anchored acetyltransferase complex, these acetylated N-terminal amino acids are recognized by the membrane-embedded E3 ubiquitin ligase, Doa10 (5). This acetylation-dependent degradation is known as the Ac/N-degron pathway. The degradation of proteins with an N-terminal proline recognized by the glucose-induced degradation-deficient (GID) ubiquitin ligase complex is called the Pro/N-degron pathway (6).

The N-degron pathways regulate several cellular processes (2). One of the most well-studied processes is the regulation of di/tri-peptide import by the Arg/N-end pathway in *S. cerevisiae*. The transcriptional expression of *PTR2*, which encodes a transmembrane di/tri-peptide transporter, is suppressed by the transcriptional repressor, Cup9 (7). When both the type-1 and type-2 substrate-binding sites of Ubr1 are occupied by dipeptides, a distinct substrate-binding site of Ubr1 recognizes the C-terminus-proximal region of Cup9 (8, 9). The degradation of Cup9 then induces *PTR2* expression and promotes di/tri-peptide uptake. This is a positive feedback mechanism for the effective uptake of di/tri-peptides as nutrient sources in response to environmental conditions (10).

The filamentous fungus, *Aspergillus oryzae*, secretes various hydrolytic enzymes for the degradation of raw materials and is used for the production of traditional Japanese fermented foods, such as sake, soy sauce, and miso (soybean paste) (11). *A. oryzae* has more peptidase genes than other closely related *Aspergillus* species, and these translational products have been used as industrial enzymes for various food-processing and pharmaceutical applications (12). On the other hand, although *A. oryzae* is used as a host for producing homologous and heterologous proteins (13, 14), degradation by peptidases is one of the major problems for efficient production (15). Therefore, controlling peptidase gene expression in *A. oryzae* is critical for various industrial applications. Although the expression of peptidase genes in filamentous fungi is regulated by several transcription factors, such as FlbC, AreA, CreA, AmdX, and XprG (16–21), the Zn(II)2Cys6-type transcription factor, PrtT, plays a central role in the regulation of extracellular peptidase gene expression (22–26). In *A. oryzae*, the expression of most extracellular peptidase genes is positively or negatively regulated by PrtR (ortholog of PrtT) in response to a nitrogen source (27). In filamentous fungi, the expression of hydrolytic genes is often regulated in concert with the expression of transporter genes that import the substrates of hydrolytic genes expression or the degradation products of polymers by hydrolytic enzymes (28, 29). We previously identified three di/tri-peptide transporter genes, *potA*, *potB*, and *potC,* in *A. oryzae* and found that the expression of *potA* and *potB* is positively regulated by PrtR (30). This suggests that the gene expression of peptidases and di/tri-peptide transporters is cooperatively regulated. The disruption of *ubrA*, the ortholog of yeast *UBR1*, reduces the expression levels of POT genes, especially *potC* (30). However, the role of UbrA in regulating peptidase gene expression remains unclear. Additionally, the N-degron pathway in filamentous fungi has not been studied in detail.

In this study, we demonstrated the involvement of UbrA in the Arg/N-degron pathway and the regulation of various peptidase genes.

## RESULTS

### Involvement of UbrA in the N-degron pathway

To investigate the involvement of *A. oryzae* UbrA in the N-degron pathway, ubiquitin-fused green fluorescent protein (Ub-X-GFP) was used as a reporter substrate to determine the N-end rule. When this fusion protein is expressed in eukaryotic cells, ubiquitin is removed by deubiquitinating enzymes, resulting in the expression of X-GFP with the desired amino acid X at the N-terminus (31, 32). The mutant Ub-G76V-GFP, in which ubiquitin is not cleaved from GFP, is degraded by the proteasome independently of the N-degron pathway (31, 32). To verify whether Ub-X-GFP is available to examine the N-degron pathway in *A. oryzae*, Ub-Met (M)-GFP, Ub-Arg (R)-GFP, and Ub-G76V-GFP were expressed in the control and *ΔubrA* strains. In the control strain, M-GFP was detected by western blotting using an anti-GFP antibody, whereas R-GFP, a typical substrate of the Arg/N-degron pathway in eukaryotes, was undetectable (Fig. S1). In contrast, R-GFP was detected equally with M-GFP in the *ΔubrA* strain, and Ub-G76V-GFP was not detected in both strains (Fig. S1). This indicates that UbrA mediates the degradation of R-GFP, depending on the N-terminal arginine. In addition, the results suggest that Ub-X-GFP can be used to examine the N-degron pathway in *A. oryzae*. Using this system, GFPs containing other 18 amino acids at the N-terminus were expressed in the control strain. In addition to R-GFP, the signals of eight other GFPs, Asp (D)-, Pro (P)-, Trp (W)-, Phe (F)-, His (H)-, Asn (N)-, Lys (K)-, and Tyr (Y)-GFP, were hardly detected (Fig. 1A). Although Glu (E)-GFP and Gln (Q)-GFP were detectable, their amounts were approximately half of that of M-GFP. The remaining eight X-GFP species were detected in similar or higher amounts than that of M-GFP. To determine whether reduced abundance X-GFPs were degraded in a UbrA-dependent manner, they were expressed in the *ΔubrA* strain. In addition to R-GFP, D-, W-, F-, H-, N-, K-, Y-, E-, and Q-GFPs, excluding P-GFP, were detected in similar amounts to that of M-GFP in the *ΔubrA* strain (Fig. 1B). These results indicated that *A. oryzae* UbrA is essential for the Arg/N-degron pathway.

**Fig. 1.**
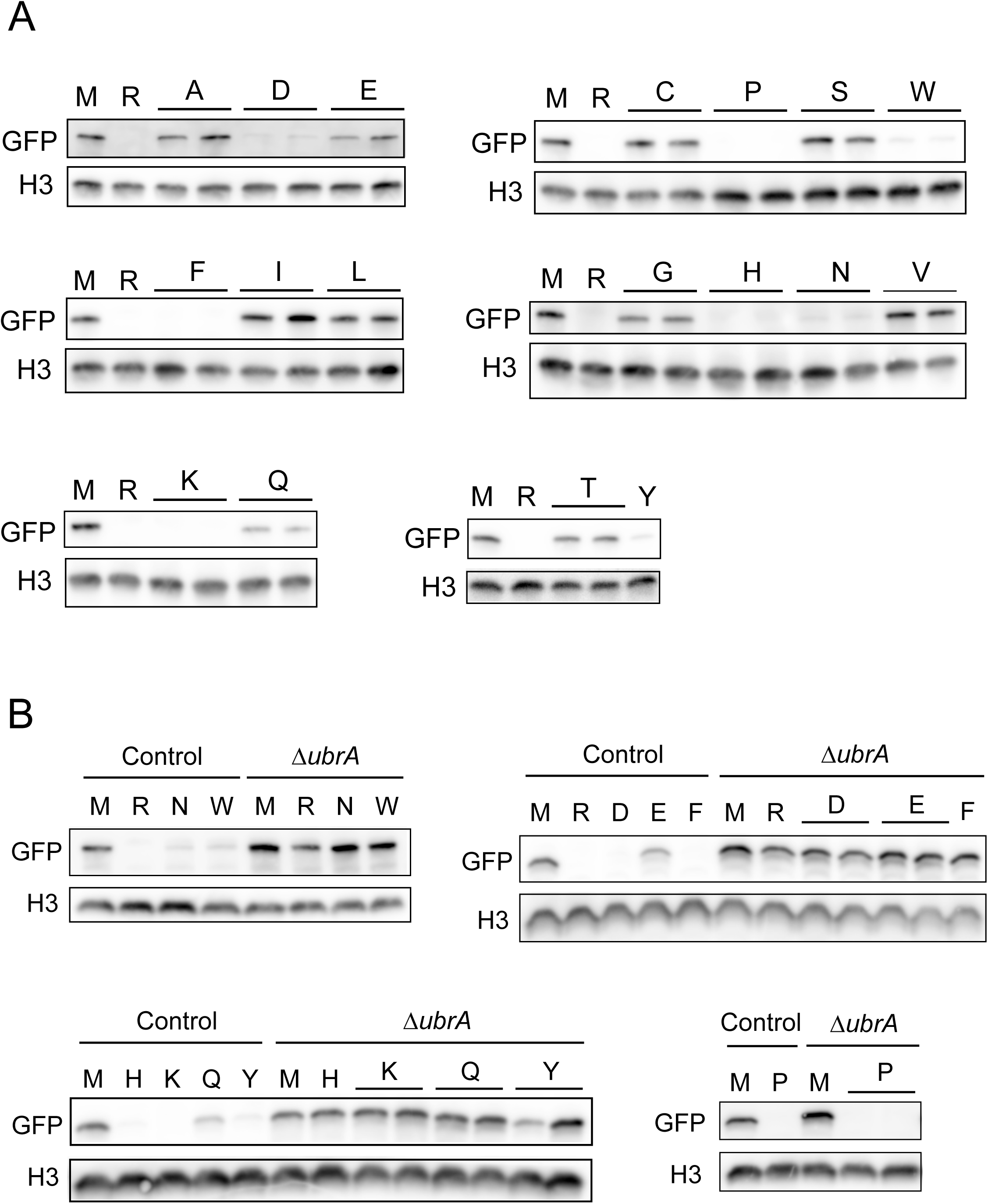
Effect of *ubrA* disruption on the abundance of X-GFPs. (A) Western blot analysis of X-GFPs in the control (A) and *ΔubrA* (B) strains. Approximately 2 × 10^7^ conidiospores of one or two Ub-X-GFP expressing strain/s were grown at 30 °C for 20 h in liquid CD + 0.1% polypepton. After harvesting the mycelium, approximately 30 μg extracted intracellular protein was subjected to western blot analysis using anti-GFP antibody. Histone H3 (H3) was detected as loading control by anti-histone H3 antibody.

### Involvement of UbrA in peptidase production in a submerged culture

The *ΔubrA* strain described above has two auxotrophic mutations that can be used as selection markers for transformation (30). Since auxotrophic mutations might affect nitrogen metabolism and peptidase expression, we regenerated the *ΔubrA* strain without auxotrophic mutation using the RIB40*ΔligDΔpyrG* strain as a host (33). To investigate the relationship between UbrA and PrtR in peptidase production, a double disruption mutant of *ubrA* and *prtR* was generated using RIB40*ΔligDΔpyrGΔprtR* strain as a host (27). A loop-out-type *pyrG* marker recycling system was used for *ubrA* disruption (Fig. S2). Following complementation with *pyrG* and *ligD* (Fig. S3), single- and double-disruption strains were used to analyze peptidase production. To examine the involvement of UbrA in peptidase production, the activities of acidic endopeptidase in the culture supernatants of these disrupted strains were compared with those of control and *ΔprtR* strains. After 48 h of cultivation in liquid medium with soy protein as the nitrogen source (CD/soy), the proteolytic activity using casein as the substrate at pH 3.0 was reduced by approximately 70% and 85% by *ubrA* and *prtR* disruption, respectively (Figs. 2A and 2B). To determine the type of peptidase in the culture supernatant, inhibitor assays were performed using pepstatin A, which specifically inhibits aspartic proteases (APases), and antipain, which inhibits serine/cysteine proteases (Figs. 2A and 2 B). The acidic endopeptidase activity of the control strain was 80% and 26% inhibited by pepstatin A and antipain, respectively. In the *ΔubrA* strain, the activity was 89% inhibited by pepstatin A and 45% by antipain. This suggests that *ubrA* disruption significantly reduces the production of both APases and serine/cysteine proteases. Although the *ΔubrA* and *ΔubrAΔprtR* strains formed compact colonies on the agar medium (Fig. S4), the semi-dry mycelia weight of the *ΔubrA* strain after cultivation in CD/soy medium was comparable to those of the control and *ΔprtR* strains (Table S1), indicating that the decrease in acidic endopeptidase activity by *ubrA* disruption was not due to differences in growth. Moreover, *ubrA* and *prtR* disruption reduced acidic carboxypeptidase activity by approximately 75% and 95%, respectively (Fig. 2C). This activity was completely abolished by the double disruption of *ubrA* and *prtR*. These results suggest that the disruption of *ubrA* reduces the production of a wide range of acidic peptidases.

**Fig. 2.**
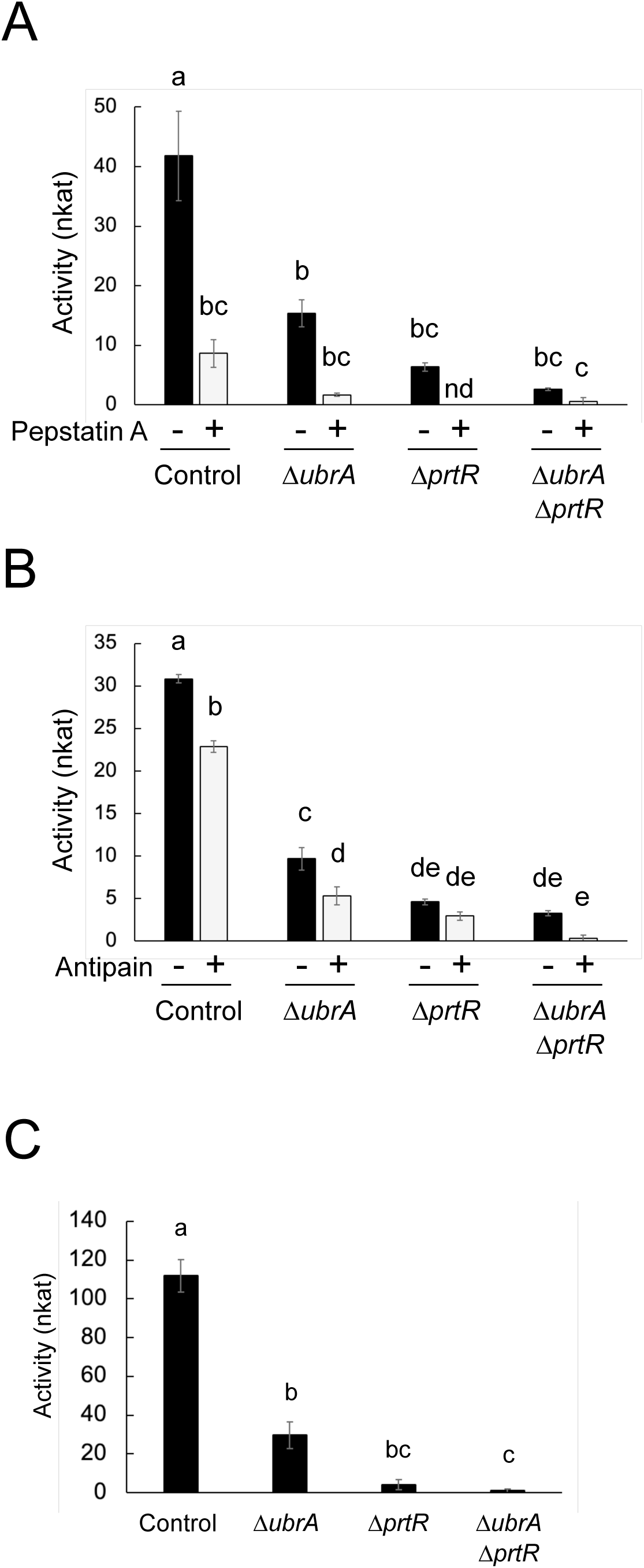
Effect of *ubrA* disruption on the acidic peptidase production in submerged culture. (A) Acid endopeptidase activity with or without pepstatin A. Approximately 2.5 × 10^7^ conidiospores of each strain were grown at 30 °C for 48 h in liquid CD/soy medium containing 0.6% soy protein as a nitrogen source. Activities of the culture supernatants were measured using 2% casein (pH 3.0) as the substrate with pepstatin A (+) or dimethyl sulfoxide (-), the solvent for pepstatin A. nd means not detected. (B) Acid endopeptidase activity with or without antipain. Activities of the culture supernatants were measured with antipain (+) or water (-). (C) Acid carboxypeptidase activity. Activities of the culture supernatants were measured using 1 mM Z-Glu-Tyr (pH 3.5) as the substrate. Error bars indicate the standard error of three biological replicates. Statistical analysis was performed using the Tukey–Kramer method. Different lowercase letters indicate significant differences (*p* < 0.05).

### Involvement of UbrA in the transcriptional expression of peptidase genes in a submerged culture

To investigate the involvement of UbrA in the transcriptional expression of peptidase genes, the mRNA levels of major peptidase genes after cultivation in liquid CD/soy medium for 36 h were examined by reverse transcription-quantitative PCR (RT-qPCR). The mRNA levels of six endopeptidase genes (*pepO*, *deuA*, *pipA*, *np3*, *aorA*, and *aorB*) were reduced following *prtR* disruption (Fig. 3A). The expression levels of all five genes, except *aorB*, were also reduced by *ubrA* disruption (Fig. 3A). Specifically, the expression levels of *pepO*, *pipA*, and *aorA* encoding APase, glutamic endopeptidase, and aorsin A, respectively were dramatically reduced by *ubrA* disruption (Fig. 3A). In contrast, the expression levels of *deuB*, *np1*, *aorB* encoding deuterolysin B, metalloendopeptidase, and aorsin B, respectively, were increased by *ubrA* disruption, although they were unchanged or reduced by *prtR* disruption (Fig. 3A). The expression levels of two carboxypeptidase genes, *ocpA* and *ocpO,* were dramatically reduced by the disruption of *ubrA* (Fig. 3B), suggesting that the decrease in carboxypeptidase activity caused by the disruption of *ubrA* is attributed to the reduced expression of these genes. The expression level of another carboxypeptidase gene, *ocpB,* was increased by *prtR* disruption but was not altered by *ubrA* disruption. These results suggest that UbrA regulates the transcription of a wide range of peptidase genes through both cooperative and independent interactions with PrtR.

**Fig. 3.**
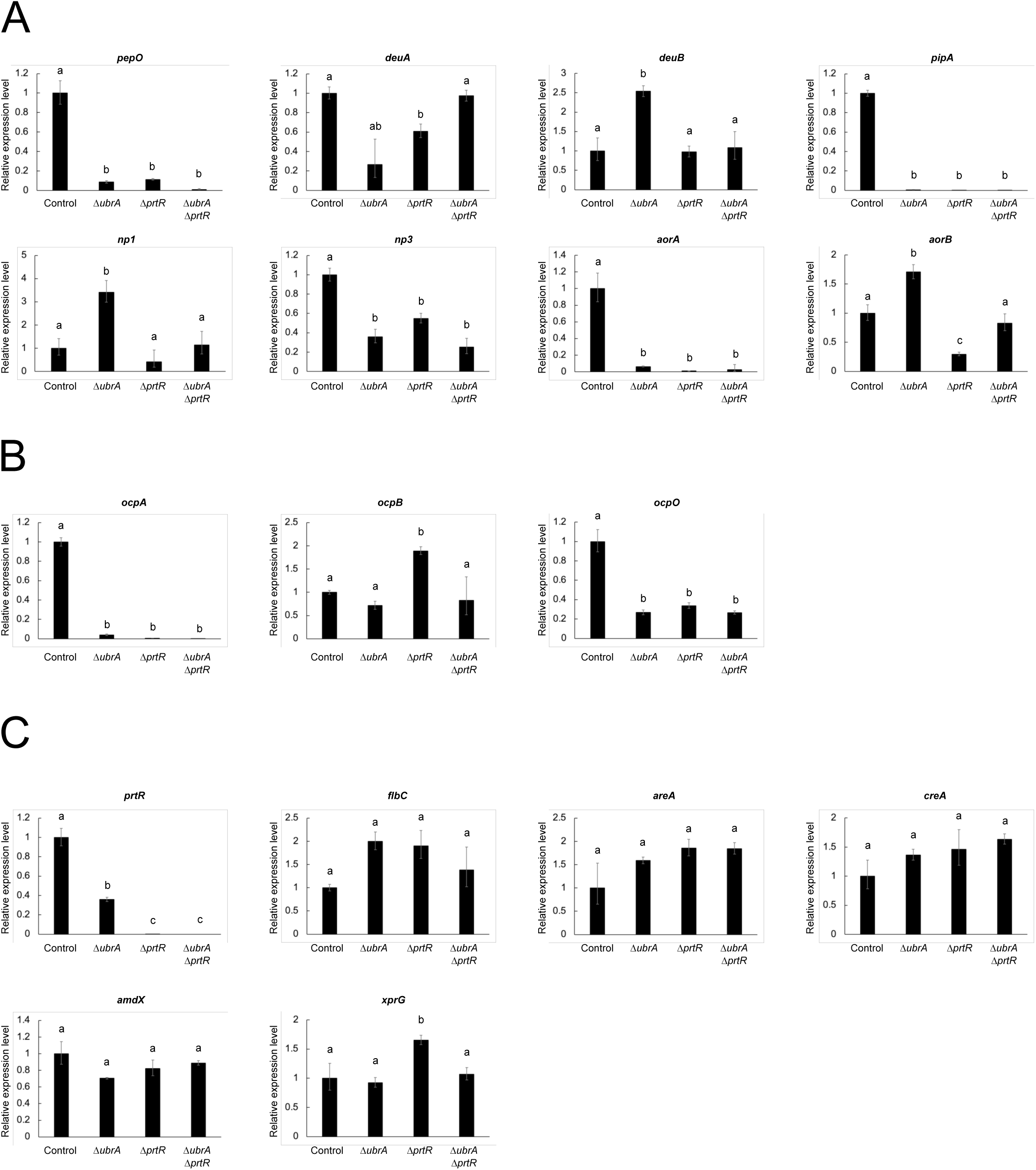
Effect of *ubrA* disruption on the transcriptional expression of peptidase genes in submerged culture. RT-qPCR analysis of endopeptidase (A), acid carboxypeptidase (B), and transcription factor genes (C) involved in the regulation of peptidase gene expression after cultivation in liquid CD/soy medium for 36 h. Error bars indicate the range of SE determined by the standard deviations using ΔΔC_T_ values of three biological replicates. Statistical analysis was performed using the Tukey–Kramer method. Different lowercase letters indicate significant differences (*p* < 0.05).

Next, we examined whether the expression levels of genes encoding transcription factors regulating peptidase gene expression were affected by *ubrA* disruption (Fig. 3C). The *prtR* expression level decreased by approximately 65% after *ubrA* disruption. The mRNA levels of the other five transcription factors, AreA, CreA, AmdX, XprG, and FlbC, were not significantly affected by either *ubrA* or *prtR* disruption. These results suggested that the disruption of *ubrA* specifically reduced *prtR* expression.

### Involvement of UbrA in the transcriptional expression of peptidase genes in a solid-state culture

*A. oryzae* produces larger amounts of hydrolytic enzymes in solid-state than in liquid cultures, particularly in solid-state culture using wheat bran as the substrate (34). To investigate the involvement of UbrA in peptidase production under solid-state cultivation, all strains were grown in solid-state culture using wheat bran as the substrate, and casein proteolytic activity at pH 3.0 in the crude enzyme extracts was measured. In the solid-state cultivation, the *ΔubrA* strain showed more vigorous conidiospore formation than the control and *ΔprtR* strains, and the growth of the *ΔubrAΔprtR* strain was slightly slower than the other strains (Fig. S5). The proteolytic activity of casein was reduced by approximately 92% and 26% by *prtR* and *ubrA* disruption, respectively (Fig. 4A). Most of the activity was abolished by the double disruption of *prtR* and *ubrA*. To investigate the effect of *ubrA* disruption on transcriptional expression, the mRNA levels of *prtR* and peptidase genes were examined (Fig. 4B). As in the submerged culture, disruption of *ubrA* reduced *prtR* expression to approximately 40% of that in the control strain. The expression levels of *pepO*, *np1*, and *ocpA* were reduced by the disruption of *ubrA*. However, their levels were only slightly reduced compared with those caused by *prtR* disruption. In addition, the expression levels of *dueA*, *deuB*, *aorA*, and *ocpO* were significantly reduced by *prtR* disruption, whereas they remained mostly unchanged by *ubrA* disruption. The expression levels of *pipA* and *ocpB* were further reduced by *ubrA* disruption than by *prtR* disruption. These results suggest that disruption of *ubrA* affects the expression of some peptidase genes in solid-state culture, but that the effect of UbrA on peptidase gene expression is greater in submerged culture than in solid-state culture.

**Fig. 4.**
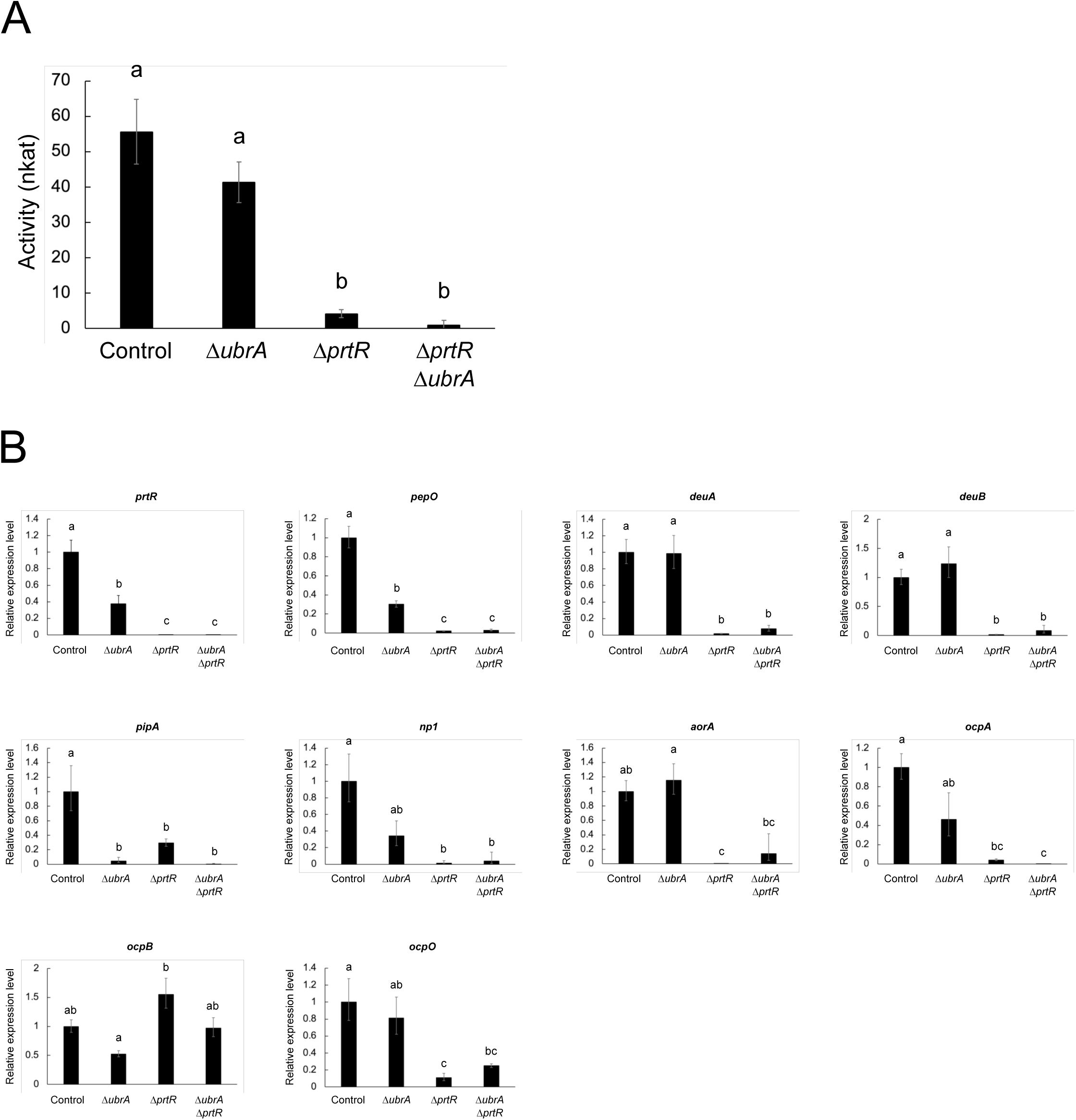
Effect of *ubrA* disruption on the expression of peptidases in solid-state culture. (A) Acid endopeptidase activity. Approximately 3 × 10^6^ conidiospores of each strain were grown at 30 °C for 36 h in wheat bran medium. Activities of the crude enzyme extracts were measured using 2% casein (pH 3.0) as the substrate. Error bars indicate the standard deviations of three independent experiments. (B) RT-qPCR analysis of *prtR* and peptidase genes. Error bars indicate the range of SE determined by the standard deviations using ΔΔC_T_ values of three biological replicates. Statistical analysis was performed using the Tukey–Kramer method. Different lowercase letters indicate significant differences (*p* < 0.05).

### Involvement of UbrA in the expression of alkaline protease and alkaline-responsive genes

*A. oryzae* produces an alkaline serine protease called oryzin (AlpA) (35, 36). To investigate the involvement of UbrA in alkaline protease production, proteolytic activity against casein at pH 9.0 was measured (Figs. 5A and 5B). In the control strain, no activity was detected when *A. oryzae* was cultured in the CD/soy liquid medium, whereas high activity was detected in the solid-state medium. When *ubrA* was disrupted, a low but distinct activity was detected when *A. oryzae* was cultured in CD/soy liquid medium, and the activity in solid-state culture was comparable to that of the control strain. The *ΔprtR* and *ΔubrAΔprtR* strains had no detectable activity in CD/soy liquid culture and only slight activity in solid-state culture. To determine whether the detectable activity in the CD/soy liquid culture was derived from oryzin, we measured the degradation activity toward Suc-Leu-Leu-Val-Tyr-MCA, which is specifically degraded by a chymotrypsin-type serine protease. Compared with the control strain, the *ΔubrA* strain showed approximately 7-fold more activity, whereas little activity was detected in the *ΔprtR* and *ΔubrAΔprtR* strains (Fig. 5C). These results suggest that UbrA is not involved in PrtR-dependent oryzin production in solid-state culture but has a role in repressing oryzin production in CD/soy liquid culture. To determine whether the induction of origin production by *ubrA* disruption depended on transcriptional expression, we examined the transcriptional expression level of *alpA*. The expression level of *alpA* increased approximately 1.5-fold in the *ΔubrA* strain compared with the control strain whereas the expression level decreased to approximately 30% in the *ΔprtR* and *ΔubrAΔprtR* strains (Fig. 5D). This suggests that UbrA is partially responsible for repressing *alpA* expression.

**Fig. 5.**
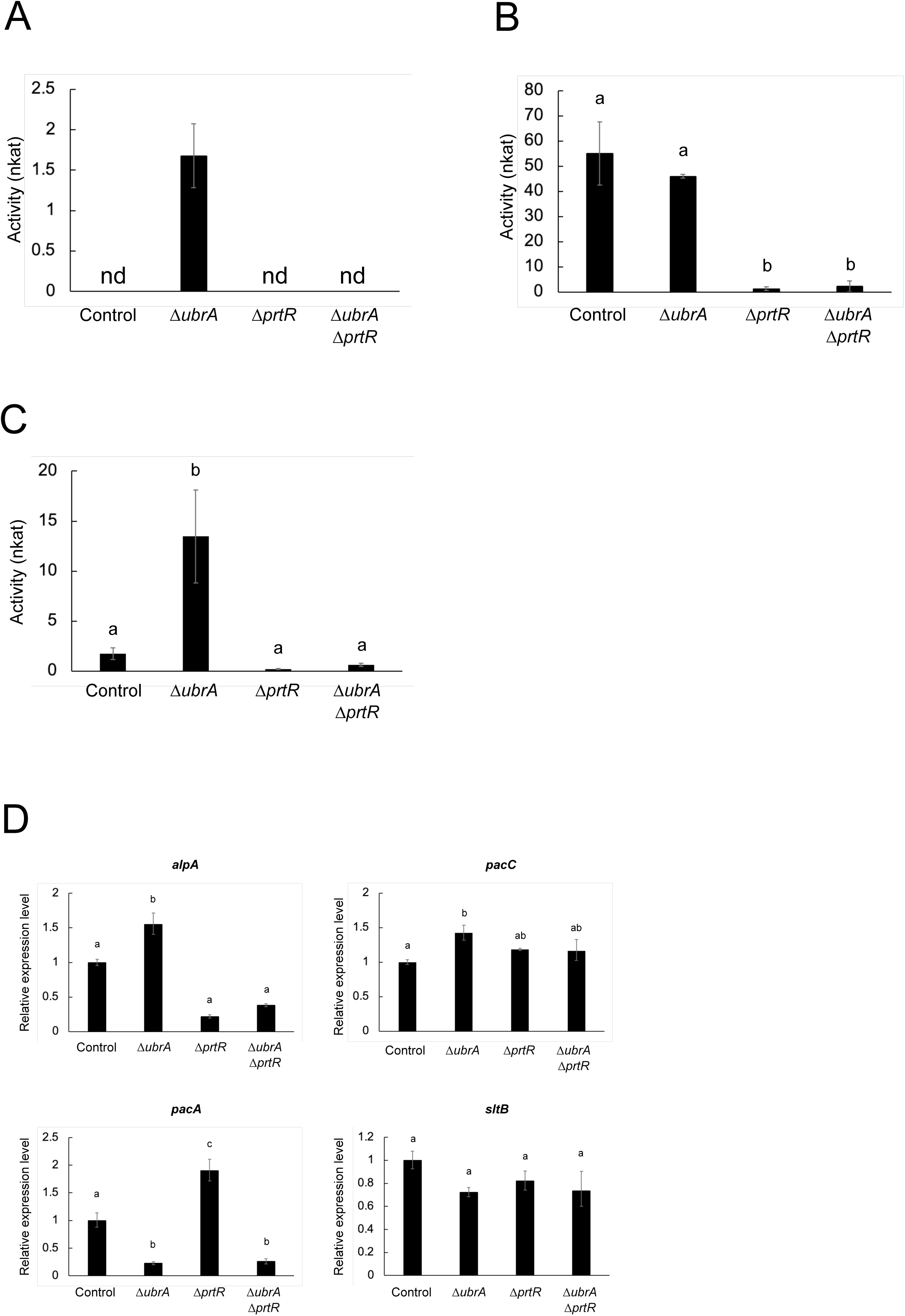
Effect of *ubrA* disruption on the alkaline peptidase expression. Alkaline endopeptidase activity in submerged (A) and solid-state (B) cultures. Activities of the culture supernatants after cultivation in CD/soy medium for 48 h and crude enzyme extracts after cultivation in wheat bran medium for 36 h were measured using 2% casein (pH 9.0) as the substrate. nd means not detected. (C) Activity of chymotrypsin-type serine protease. Activities of the culture supernatants were measured using Suc-Leu-Leu-Val-Tyr-MCA (pH 10.0) as the substrate. (D) RT-qPCR analysis of *alpA* and alkaline-responsible factor genes. Error bars indicate the range of SE determined by the standard deviations using ΔΔC_T_ values of three biological replicates. Statistical analysis was performed using the Tukey– Kramer method. Different lowercase letters indicate significant differences (*p* < 0.05).

Although PrtR is required to induce *alpA* expression (22), pH-responsive transcription factor PacC also regulates *alpA* expression (37). Therefore, we examined the expression levels of *pacC* and other alkaline-responsive factors (Fig. 5D). The expression level of *pacC* increased approximately 1.5-fold by disruption of *ubrA*. In contrast, the expression level of the acidic phosphatase gene, *pacA*, in which expression is repressed by PacC (37), was significantly reduced by *ubrA* disruption. The expression levels of *sltB*, which is involved in the expression of cation/alkaline stress response genes and is induced by cation/alkaline stress (38, 39), were not increased by *ubrA* disruption. Moreover, the pH of the culture supernatant of each strain after cultivation in the CD/soy medium was approximately 4.0 (Table S2). These results suggest that disruption of *ubrA* induces the expression of *alpA* through PacC, independent of pH.

### Involvement of UbrA in the gene expression of dipeptidyl and tripeptidyl peptidases

We previously identified three di/tri-peptide transporter genes in *A. oryzae*. *ubrA* disruption reduced *potA* and *potC* expression levels but not *potB* when glycine or leucylglycine was used as the nitrogen source (30). Consistently, the expression levels of *potA* and *potC*, but not *potB*, were reduced by the disruption of *ubrA* when cultured in CD/soy medium (Fig. 6A). Di/tri-peptides are generated from polypeptides through cleavage by dipeptidyl peptidase (DPP) and tripeptidyl peptidase (TPP); *A. oryzae* produces three DPP and TPP (40, 41, 42). To investigate the involvement of UbrA in the generation of di/tri-peptides, the expression levels of *dpp* and *tpp* were examined (Figs. 6B and 6C). The expression levels of *dppB* encoding Xaa-Prolyl DPP as well as the two TPP genes, *tppA* (*sedB*) and *tppB,* were significantly reduced by *ubrA* disruption. In addition, TPP activities against Phe-Pro-Ala-pNA and Ala-Ala-Phe-pNA substrates were significantly reduced by the disruption of *ubrA* and *prtR* (Fig. 6D). Although the disruption of *prtR* also significantly reduced the expression levels of *potA* and *tppB*, the expression level of *tppA* was more reduced by the disruption of *ubrA* compared with the disruption of *prtR*. Moreover, *prtR* disruption did not affect the expression of *potC* or *dppB*. These results suggest that UbrA regulates DPP and TPP gene expression, and not only of di/tri-peptide transporter genes, independent of PrtR.

**Fig. 6.**
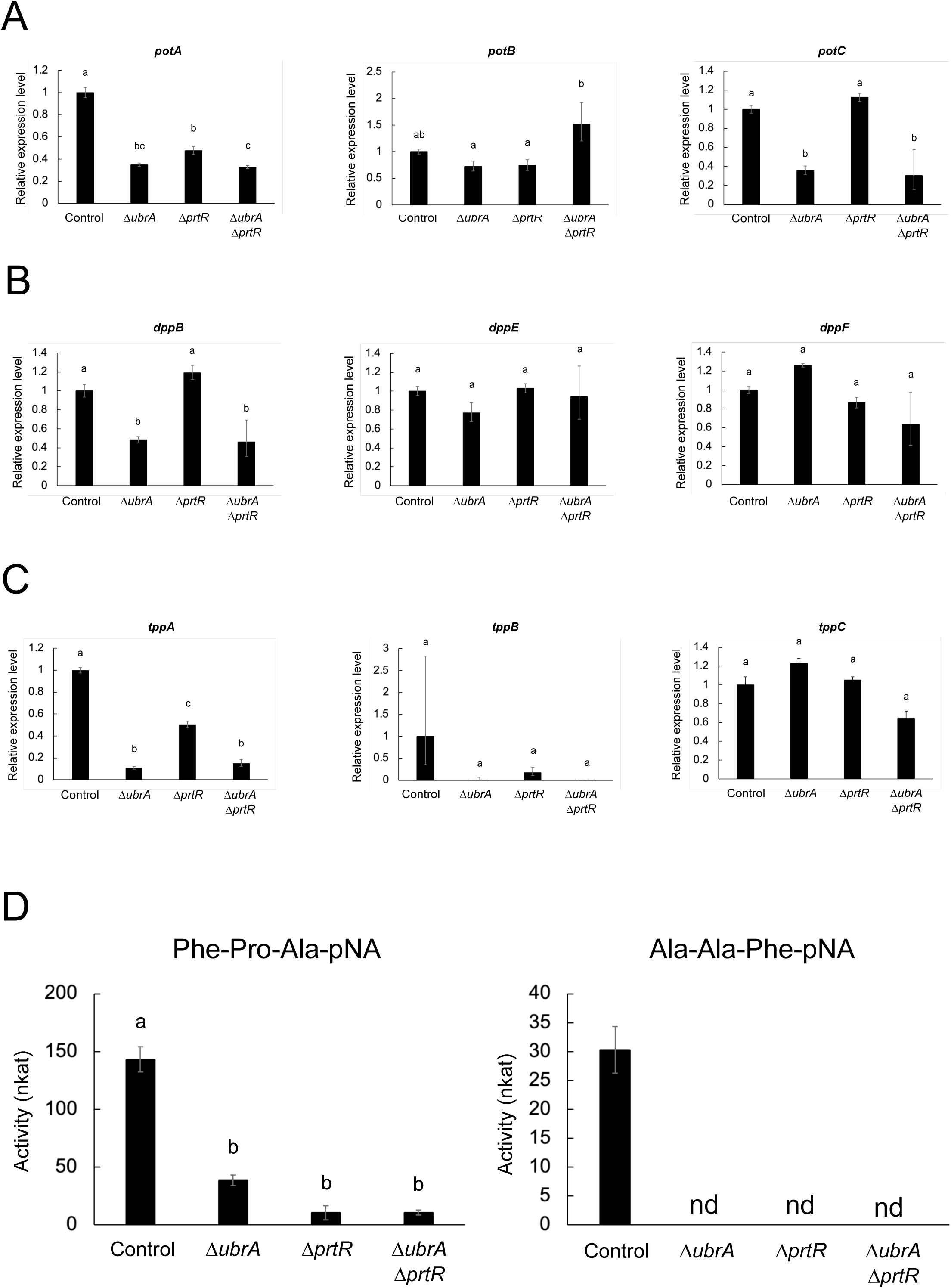
Effect of *ubrA* disruption on the expression of dipeptidyl- and tripeptidyl-peptidase genes. RT-qPCR analysis of di/tri-peptide transporter (A), dipeptidyl peptidase (B), and tripeptidyl peptidase (C) genes after cultivation in CD/soy medium for 36 h. Error bars indicate the range of SE determined by the standard deviations using ΔΔC_T_ values of three biological replicates. (D) Tripeptidyl peptidase activity. Activities of the culture supernatants were measured using Phe-Pro-Ala-pNA and Ala-Ala-Phe-pNA (pH 5.5) as the substrates. Error bars indicate the standard deviations of three independent experiments. Statistical analysis was performed using the Tukey–Kramer method. Different lowercase letters indicate significant differences (*p* < 0.05). nd means not detected.

## DISCUSSION

The Arg/N-degron pathway regulates various cellular functions (2). Notably, it is important for obtaining nitrogen sources in budding yeast because it regulates the expression of di/tri-peptide transporter gene. However, its involvement in regulating nitrogen metabolism, other than di/tri-peptide import in microorganisms, is unknown. In this study, we showed that UbrA regulates peptidase gene expression in *A. oryzae*. This suggests that the Arg/N-degron pathway broadly regulates the nitrogen metabolism in filamentous fungi.

Although the stabilization of R-GFP by disrupting the *UBR1* ortholog in *Fusarium verticillioides* has been reported (43), there is no detailed information on the N-degron pathways in filamentous fungi. In this study, we revealed that GFPs with N-terminal amino acids recognized by *S. cerevisiae* Ubr1, except for leucine and isoleucine, were degraded in a UbrA-dependent manner in *A. oryzae* (Figs. 1A and 1 B). This suggests that the Arg/N-degron pathways in *S. cerevisiae* and filamentous fungi are similar. GFP with N-terminal proline was not detected even in the *ΔubrA* strain (Fig. 1B). In *S. cerevisiae*, several glycogenic enzymes are cleaved by MetAP, leading to a proline at the N-terminus, which is recognized by the GID ubiquitin ligase complex and resulting in ubiquitination and proteasome-dependent degradation in the presence of glucose (6). Since the Pro/N-degron pathway is highly conserved from yeast to humans (44), GFP with an N-terminal proline in *A. oryzae* was probably degraded in a GID ubiquitin ligase complex-dependent manner. Although GFPs with N-terminal leucine and isoleucine were not reduced in the control strain, whether UbrA recognizes these N-terminal amino acids requires further investigation.

Although disruption of *ubrA* reduces *prtR* expression, the molecular mechanism by which UbrA regulates peptidase expression is unknown. In *S. cerevisiae*, the binding of dipeptides to the UBR-box sites of Ubr1 allows the recognition of the transcriptional repressor, Cup9, at a distinct substrate-binding site of Ubr1, which induces the expression of *PTR2* encoding a di/tri-peptide transporter by promoting the degradation of Cup9 (7–10). Therefore, one hypothesis is that Ubr1 degrades transcriptional repressors to suppress *prtR* and peptidase gene expression. In filamentous fungi, NmrA is a corepressor of the GATA transcription factor, AreA, which regulates the expression of nitrogen metabolism genes under nitrogen limitation and starvation conditions (45, 46). It has been reported that NmrA is cleaved by three proteases, including trypsin-like serine protease, PrmB, when AreA function is required (47, 48). Although the relationship between PrtR and NmrA remains unknown, UbrA may degrade intact NmrA or its cleaved products. Identifying the proteins recognized by UbrA is expected to provide novel insights into the regulatory mechanisms of peptidase gene expression.

Unlike several genes encoding acidic and neutral peptidases, the expression of *alpA* in submerged culture was induced by *ubrA* disruption (Fig. 5D). Because *ubrA* disruption also increased *pacC* expression and markedly decreased the *pacA* expression (Fig. 5D), this induction of *alpA* expression is likely to be mediated by PacC. It is well known that PacC is activated after truncation mediated by the Pal pathway (49–53). In *A. nidulans*, the sensing of alkaline pH by the transmembrane protein, PalH, induces post-translational modification of the arrestin-related trafficking adaptor protein, PalF, followed by the recruitment of PalA to intact PacC (PacC^72^). The calpain-like protease, PalB, binds to the PalA-PacC complex and cleaves PacC. The C-terminus of the resultant N-terminus of PacC (PacC^53^) was further truncated by the proteasome to generate the active form of PacC (PacC^27^). Thus, intact or truncated PacC may be degraded in a UbrA-dependent manner. Because the pH of the culture supernatant of each strain used in this study after cultivation in CD/soy medium was approximately 4.0 (Table S2), we speculated that UbrA plays a role in the clearance of active PacC under conditions where PacC is nonfunctional. Therefore, it is necessary to examine whether intact or truncated PacC is subject to degradation in a UbrA-dependent manner.

In *A. oryzae*, the expression of some hydrolytic genes is induced specifically in solid-state culture, and this specific expression requires FlbC (16). This indicates that the regulatory mechanisms for hydrolytic enzyme gene expression in liquid and solid-state cultures are very different and are more complex in solid-state cultures. The effect of *ubrA* disruption on peptidase gene expression was more limited in solid-state culture than in submerged culture (Fig. 4B). This suggests that the contribution of UbrA to the regulation of complex gene expression in solid-state culture was not significant. In addition, *ubrA* disruption promoted the formation of conidiospores in solid-state cultures (Fig. S5); therefore, the effect of this morphological change on the regulation of peptidase gene expression should be considered.

The regulation of *PTR2* expression through the degradation of Cup9 is one of the most well-studied functions of Ubr1. This is a positive feedback mechanism in which *PTR2* expression is induced by dipeptides imported through Ptr2 (10). Similar to *PTR2* in budding yeast, the expression of *potA* and *potC* in *A. oryzae* is regulated in a UbrA-dependent manner (Fig. 6A). In this study, we showed that UbrA is also involved in the expression of DPP and TPP genes in *A. oryzae* (Figs. 6B and 6C). UbrA-dependent expression of the DPP and TPP genes likely facilitates the positive feedback of di/tri-tripeptide transporter gene expression in an environment where proteins are available as a nutrient source. Because the Cup9 ortholog is not conserved in the genomes of filamentous fungi, the identification of transcriptional repressors of the POT, DPP, and TPP genes is necessary to understand the di/tri-peptide acquisition mechanism in filamentous fungi.

The greater effect of *ubrA* disruption than of *prtR* disruption on the expression of genes encoding DPP, TPP, and POT suggests that UbrA-mediated regulation is independent of PrtR. In this study, the expression levels of *tppC*, *potB*, *potC*, and all three DPP genes were unaffected by *prtR* disruption (Figs. 6A–C). In contrast, when *A. oryzae* was cultured in CD using casein as the nitrogen source, the expression of all three *dpp* genes was increased by *prtR* disruption (27). In *A. fumigatus*, *prtT* deletion reduced the expression of *dppIV* and *sedB* (*tppA*) but did not alter the expression of *dppV* when cultured in liquid medium containing BSA (24). In *Penicillium oxalicum*, *prtT* deletion enhances *dppV* expression when cultured in a liquid medium containing casein (54). These results suggest that PrtR (PrtT)-dependent gene expression varies with the culture conditions. As Ubr1 is activated by the binding of dipeptides, UbrA activation is also expected to be greatly affected by the peptides present in the medium. Therefore, the effects of *ubrA* disruption on peptidase gene expression under various culture conditions should be investigated in future studies.

In conclusion, this study revealed that *A. oryzae* UbrA is essential for the Arg/N-degron pathway, positively regulates major acid peptidase genes, and negatively regulates alkaline peptidase gene. This study also revealed that UbrA regulates DPP and TPP gene expression in concert with the expression of di/tri-peptide transporter genes. These results suggest that UbrA regulates the expression of various peptidase genes to facilitate positive feedback of di/tri-tripeptide transporter genes. Further understanding of UbrA function will provide important information regarding nitrogen metabolism in filamentous fungi.

## MATERIALS AND METHODS

### Strains and media

The *ΔligD*::*ptrA* (55) and *ΔubrA* (30) strains were used as recipient strains to express Ub-X-GFPs. RIB40*ΔligDΔpyrG* (33) and RIB40*ΔligDΔprtRΔpyrG* (27) strains were used as host strains for *ubrA* disruption to examine peptidase expression. Deletion-control (33) and RIB40*ΔprtR* (27) strains were used as control and *ΔprtR* strains, respectively, to examine peptidase expression. The *A. oryzae* strains used in this study are listed in Table S3. *Escherichia coli* DH5α was used to construct and propagate the plasmids. Czapek–Dox (CD) medium containing 0.6% NaNO_3_, 0.05% KCl, 0.2% KH_2_PO_4_, 0.05% MgSO_4_, trace amounts of FeSO_4_, ZnSO_4_, CuSO_4_, MnSO_4_, Na_2_B_4_O_7_, (NH_4_)_6_Mo_7_O_24_, and 1% glucose was used as the standard minimal medium for *A. oryzae* cultivation. L-Methionine was supplemented at a final concentration of 0.003% to cultivate the *sC*-deficient strain. HIPOLYPEPTON N (Nihon Pharmaceutical Co., Ltd., Tokyo, Japan) was added to the liquid CD medium at a final concentration of 0.1% to promote growth when examining the N-end rule. Soy protein (0.6 %; Fuji Oil Co., Ltd., Osaka, Japan) was used as the nitrogen source for analysis of peptidase production. To examine peptidase production in solid-state culture, wheat bran medium was prepared by adding 6 mL reverse osmosis water to 4.2 g wheat bran (Showa Sangyo Co., Ltd., Tokyo, Japan).

### DNA fragments for expression of Ub-X-GFPs

Plasmid DNA for the expression of Ub-M-GFP, Ub-R-GFP, and Ub-G76V-GFP were constructed as follows: the DNA fragments encoding Ub-M-GFP, Ub-R-GFP, and Ub-G76V-GFP were amplified through PCR using the templates (pYES2-Ub-M-GFP, pYES2-Ub-R-GFP, and YES2-Ub-G76V-GFP, respectively) (32) purchased from Addgene (Addgene plasmids #11952, 11953, and 11954) and the primers Ub-GFPsen MTS382 and Ub-GFPanti MTS383. The amplified DNA fragments were introduced between the *thiA* promoter and *agdA* terminator of *Not*I-digested pNthiA (56) using the SLiCE method (57). The resultant plasmids were designated as pNT-Ub-M-GFP, pNT-Ub-R-GFP, and pNT-Ub-G76V-GFP. These plasmids were digested with *Hpa*I and integrated at the *niaD* locus via a single crossover in the *ΔligD*::*ptrA* and *ΔubrA* strains.

To analyze the Arg/N-degron pathway, DNA fragments for the expression of Ub-X-GFP were constructed as follows: DNA fragments of the partial *niaD* marker and the Ub-M-GFP or Ub-R-GFP expression cassette regions were amplified through PCR using pNT-Ub-M-GFP or pNT-Ub-R-GFP as templates and the primer sets niaDantiIFsC MTS471 + TagdA-niaDanti MTT576 and niaD5-TagdAanti MTT577 + niaD3-PthiAsen MTT578. The downstream region of *niaD* was amplified through PCR using the genomic DNA of *A. oryzae* RIB40 strain as a template and the primers PthiA-niaD3sen MTT579 and niaD3anti MTT580. These three PCR fragments were mixed, and a second round of PCR was performed with the primers niaDantiIFsC MTS471 and niaD3anti MTT580. The resulting PCR fragments were integrated at the *niaD* locus via a double crossover in the *ΔligD*::*ptrA* and *ΔubrA* strains. DNA fragments for the expression of other Ub-X-GFPs were amplified through fusion PCR. Two PCR fragments were amplified from the genomic DNA of the Ub-M-GFP-expressing strain using the primer sets niaDantiIFsC MTS471 + Ub-X-GFPanti and Ub-X-GFPsen + niaD3-PthiAsen MTT578 (X indicates the amino acid to be substituted with methionine). These two PCR fragments were mixed, and a second round of PCR was performed with the primers niaDantiIFsC MTS471 and niaD3anti MTT580. The resulting PCR fragments were integrated at the *niaD* locus via a double crossover in the *ΔligD*::*ptrA* and *ΔubrA* strains. The nucleotide sequences of all primers used in this study are presented in Table S4.

### Construction of *ubrA* disruption strain used for examining peptidase production

The DNA fragment used for *ubrA* disruption was constructed as follows: the DNA fragments of *ubrA* 5ʹ flanking and 3ʹ coding regions for homologous recombination to genomic DNA of host strains were amplified from *A. oryzae* genomic DNA via PCR with primer sets ubrAup_F1 + ubrAup_R2 and ubrAdown_F5 + ubrAdown_R6. Another *ubrA* 3ʹ coding region for loop-out after *ubrA* disruption was amplified from *A. oryzae* genomic DNA through PCR with primer sets ubrAdown_loopF3 and ubrAdown_loopR4. The *A. nidulans pyrG* marker fragment was amplified from the pUC/pyrG/niaD plasmid (58) through PCR using the primer sets AnpyrGsen and AnpyrGantiPstI. The four resulting amplified fragments were introduced into linearized pUC19 (Takara Bio Inc., Shiga, Japan) using NEBuilder HiFi DNA Assembly Master Mix (New England Biolabs Japan Inc., Tokyo, Japan). The resultant plasmid, pUC/ΔubrA/pyrGloop-out, was linearized via *Sma*I digestion and introduced into *A. oryzae* protoplast.

### *A. oryzae* transformation

*A. oryzae* was transformed via protoplast polyethylene glycol method described by Gomi et al. (59). Yatalase (Ozeki Co., Hyogo, Japan) was used for the protoplast preparation.

### *pyrG* marker recycling

Selection of *A. oryzae* strains in which the *pyrG* marker was removed via loopout was conducted according to Maruyama et al. (60). The *ligD* and *pyrG* were complemented in *pyrG* removed strains, as previously described (33).

### Southern blot analysis

Extraction of *A. oryzae* genomic DNA and Southern blotting were performed as previously described (33). The probe used to confirm *ubrA* disruption was amplified via PCR using genomic DNA of *A. oryzae* RIB40 strain as a template and the primers ubrAprobeF15 and ubrAprobeR16.

### Intracellular protein extraction for GFP detection

After culturing in liquid CD + 0.1% HIPOLYPEPTON N for 20 h, mycelia were harvested using Miracloth (Merck KGaA, Darmstadt, Germany) and washed with distilled water. Mycelia were ground to a fine powder in liquid nitrogen using a mortar and pestle, suspended in 1 × Laemmli sample buffer (61), and boiled for 3 min. After centrifugation at 20,400 × *g* for 10 min, the supernatant was collected as an intracellular protein extraction sample. Protein concentrations were quantified using Qubit 4 Fluorometer and Qubit™ protein assay kit (Thermo Fisher Scientific Inc., Waltham, MA, USA). Approximately 30 μg protein was subjected to western blot analysis.

### Western blot analysis

The extracted intracellular proteins were separated by sodium dodecyl sulfate-polyacrylamide gel electrophoresis using AnykD™ Mini-PROTEAN® TGX™ gel (Bio-Rad Laboratories Inc., Hercules, CA, USA). The proteins were then transferred to a polyvinylidene difluoride membrane using the Trans-Blot® Turbo™ Transfer System (Bio-Rad Laboratories Inc.). Horseradish peroxidase-conjugated anti-GFP (B-2) (Santa Cruz Biotechnology Inc., Dallas, TX, USA), anti-Histone H3 (ab21054) (Abcam plc., Cambridge, UK), and anti-PGK1 (14) (Santa Cruz Biotechnology Inc.) antibodies were used for protein detection according to the manufacturer’s instructions. ImmunoStar® LD (FUJIFILM Wako Pure Chemical Corporation, Osaka, Japan) and Chemi-Lumi One L (Nacalai Tesque, Inc., Kyoto, Japan) were used as chemiluminescence reagents. ImageQuant™ LAS 500 (Cytiva, Tokyo, Japan) was used to detect chemiluminescent signals.

### Measurement of enzymatic activities

The activities of acidic endopeptidases and carboxypeptidases after cultivation in CD/soy medium were measured as previously described (27). To measure enzymatic activity in solid-state culture, approximately 3 × 10^6^ conidiospores of each strain were grown on wheat bran medium at 30 °C for 36 h. After cultivation, molded wheat bran culture was suspended in 60 mL deionized water and shaken for 180 min at 20 °C to extract the secreted enzymatic proteins. Subsequently, the crude enzyme extract was obtained via filtration through Miracloth and dialyzed against a 10 mM acetate buffer (pH 5.0) at 4 °C. The dialyzed crude enzyme extract was centrifuged at 15,000 × *g* and 4 °C for 30 min and subjected to measurement of enzymatic activities.

The activity of alkaline endopeptidase was measured in 0.1 M Tris-HCl buffer (pH 9.0) with casein as the substrate. The specific activity of chymotrypsin-type serine protease was measured in 0.1 M boric acid buffer (pH 10.0) with Suc-Leu-Leu-Val-Tyr-MCA as the substrate. The amount of 7-amino-4-methylcoumarin (AMC) from Suc-Leu-Leu-Val-Tyr-MCA was determined as follows: 95 µL enzyme solution diluted with 1.3 mL 0.1 M boric acid buffer (pH 10.0) was preincubated at 30 °C for 10 min, and then 5 µL 10 mM Suc-Leu-Leu-Val-Tyr-MCA preincubated at 30 °C was added to the enzyme solution. The increase in emission at 440 nm with excitation at 360 nm was monitored every 10 s at 30 °C using the Fluorescence Spectrophotometer F-2500 (Hitachi High-Technologies Corp., Tokyo, Japan). One katal of the chymotrypsin-type serine protease is defined as the enzyme amount that yields an equivalent of 1 mol AMC per second, using Z-Arg-Arg-MCA as the substrate under the conditions described above.

The activity of TPP was measured in 0.1 M citrate buffer (pH 5.5) with Phe-Pro-Ala-pNA and Ala-Ala-Phe-pNA as substrates. The amount of *para*-nitroaniline (pNA) from the substrates was determined as follows: 20 µL enzyme solution diluted with 178 µL 0.1 M citrate buffer (pH 5.5) was preincubated at 30 °C for 10 min, and then 2 µL 10 mM Phe-Pro-Ala-pNA or Ala-Ala-Phe-pNA preincubated at 30 °C was added to the enzyme solution. After 30 min of incubation at 30 °C, the absorbance of the mixture at 405 nm was measured using the SpectraMax®iD5 (Molecular Devices Japan, LLC., Tokyo, Japan). One katal of TPP is defined as the amount of enzyme that yields an equivalent of 1 mol pNA per second, using a standard curve obtained with known amounts of pNA.

### Total RNA extraction, complementary DNA synthesis, and RT-qPCR analysis

Total RNA was extracted as previously described (27, 33). Complementary DNA was synthesized using the PrimeScript FAST RT reagent kit with gDNA Eraser (Perfect Real Time) (Takara Bio Inc.). RT-qPCR analysis was performed using Mx3000P (Agilent Technologies Japan, Ltd., Tokyo, Japan). The actin gene (AO090701000065; *actA*) was used as a reference, and mRNA expression relative to the control strain was calculated using the comparative C_T_ (ΔΔC_T_) method. The standard deviation was calculated according to the ABI PRISM® 7700 Sequence Detection System User Bulletin #2 (Thermo Fisher Scientific).

## ACKNOWLEDGMENTS

This study was partially supported by the Noda Institute for Scientific Research Grant, MAYEKAWA HOUONKAI FOUNDATION Grant, Institute for Fermentation, Osaka (IFO) Grant, and MEXT/JSPS KAKENHI Grant Number 17H07001. We would like to thank Editage (www.editage.jp) for English language editing.

## CRediT authorship contribution statement

Waka Muromachi: Investigation. Mao Ohba: Investigation. Yasuaki Kawarasaki: Writing - Review & Editing. Youhei Yamagata: Conceptualization, Writing - Review & Editing. Mizuki Tanaka: Conceptualization, Investigation, Writing - Original Draft, Funding acquisition, Project administration.

## REFERENCES

1. Varshavsky A. 2019. N-degron and C-degron pathways of protein degradation. Proc Natl Acad Sci U S A 116: 358–366.

2. Varshavsky A. 2024. N-degron pathways. Proc Natl Acad Sci U S A 121: e2408697121.

3. Pan M, Zheng Q, Wang T, Liang L, Mao J, Zuo C, Ding R, Ai H, Xie Y, Si D, Yu Y, Liu L, Zhao M. 2021. Structural insights into Ubr1-mediated N-degron polyubiquitination. Nature 600: 334–338.

4. Sherpa D, Chrustowicz J, Schulman BA. 2022. How the ends signal the end: Regulation by E3 ubiquitin ligases recognizing protein termini. Mol Cell 82: 1424–1438.

5. Hwang CS, Shemorry A, Varshavsky A. 2010. N-terminal acetylation of cellular proteins creates specific degradation signals. Science 327: 973–977.

6. Chen SJ, Wu X, Wadas B, Oh JH, Varshavsky A. 2017. An N-end rule pathway that recognizes proline and destroys gluconeogenic enzymes. Science 355: eaal3655.

7. Byrd C, Turner GC, Varshavsky A.1998. The N-end rule pathway controls the import of peptides through degradation of a transcriptional repressor. EMBO J. 17: 269–77.

8. Du F, Navarro-Garcia F, Xia Z, Tasaki T, Varshavsky A. 2002. Pairs of dipeptides synergistically activate the binding of substrate by ubiquitin ligase through dissociation of its autoinhibitory domain. Proc Natl Acad Sci U S A 99:14110–14115.

9. Xia Z, Webster A, Du F, Piatkov K, Ghislain M, Varshavsky A. 2008. Substrate-binding sites of UBR1, the ubiquitin ligase of the N-end rule pathway. J Biol Chem 283: 24011–24028.

10. Hershko A, Ciechanover A, Varshavsky A. 2000. The ubiquitin system. Nat Med 6: 1073–1081.

11. Machida M, Yamada O, Gomi K. 2008. Genomics of *Aspergillus oryzae*: Learning from the History of Koji Mold and Exploration of Its Future. DNA Res 15:173– 183.

12. Ichishima E. 2016. Development of enzyme technology for *Aspergillus oryzae*, *A. sojae*, and *A. luchuensis*, the national microorganisms of Japan. Biosci Biotechnol Biochem 80: 1681–1692.

13. Christensen T, Woeldike H, Boel E, Mortensen SB, Hjortshoej K, Thim L, Hansen MT. 1998. High Level Expression of Recombinant Genes in *Aspergillus oryzae*. Nat Biotechnol 6, 1419–1422.

14. Fleissner A, Dersch P. 2010. Expression and export: recombinant protein production systems for *Aspergillus*. Appl Microbiol Biotechnol. 87: 1255–1270.

15. Jin FJ, Hu S, Wang BT, Jin L. 2021. Advances in Genetic Engineering Technology and Its Application in the Industrial Fungus *Aspergillus oryzae*. Front Microbiol. 12: 644404.

16. Tanaka M, Yoshimura M, Ogawa M, Koyama Y, Shintani T, Gomi K. 2016. The C_2_H_2_-type transcription factor, FlbC, is involved in the transcriptional regulation of *Aspergillus oryzae* glucoamylase and protease genes specifically expressed in solid-state culture. Appl Microbiol Biotechnol 100: 5859–5868.

17. Katz ME, Gray KA, Cheetham BF. 2006. The *Aspergillus nidulans xprG* (*phoG*) gene encodes a putative transcriptional activator involved in the response to nutrient limitation. Fungal Genet Biol 43: 190–199.

18. Qian Y, Sun Y, Zhong L, Sun N, Sheng Y, Qu Y, Zhong Y. 2019. The GATA-Type Transcriptional Factor Are1 Modulates the Expression of Extracellular Proteases and Cellulases in *Trichoderma reesei*. Int J Mol Sci 20: 4100.

19. Katz ME, Bernardo SM, Cheetham BF. 2008. The interaction of induction, repression and starvation in the regulation of extracellular proteases in *Aspergillus nidulans*: evidence for a role for CreA in the response to carbon starvation. Curr Genet 54: 47–55.

20. Shemesh E, Hanf B, Hagag S, Attias S, Shadkchan Y, Fichtman B, Harel A, Krüger T, Brakhage AA, Kniemeyer O, Osherov N. 2017. Phenotypic and Proteomic Analysis of the *Aspergillus fumigatus* Δ*PrtT*, Δ*XprG* and Δ*XprG*/Δ*PrtT* Protease-Deficient Mutants. Front Microbiol 8: 2490.

21. Hatamoto O, Umitsuki G, Machida M, Sano M, Tanaka A, Oka C, Maeda H, Tainaka H, Ito T, Uchikawa T, Masuda T, Matsushima K. 2010. Recombinant vector capable of increasing secretion of Koji mold protease. U.S. patent US7842799 B2.

22. Punt PJ, Schuren FH, Lehmbeck J, Christensen T, Hjort C, van den Hondel CA. 2008. Characterization of the *Aspergillus niger prtT*, a unique regulator of extracellular protease encoding genes. Fungal Genet Biol 45: 1591–1599.

23. Sharon H, Hagag S, Osherov N. 2009. Transcription factor PrtT controls expression of multiple secreted proteases in the human pathogenic mold *Aspergillus fumigatus*. Infect Immun 77: 4051–4060.

24. Bergmann A, Hartmann T, Cairns T, Bignell EM, Krappmann S. 2009. A regulator of *Aspergillus fumigatus* extracellular proteolytic activity is dispensable for virulence. Infect Immun 77: 4041–4050.

25. Hagag S, Kubitschek-Barreira P, Neves GW, Amar D, Nierman W, Shalit I, Shamir R, Lopes-Bezerra L, Osherov N. 2012. Transcriptional and proteomic analysis of the *Aspergillus fumigatus ΔprtT* protease-deficient mutant. PLoS One 7: e33604.

26. Huang L, Dong L, Wang B, Pan L. 2020. The transcription factor PrtT and its target protease profiles in *Aspergillus niger* are negatively regulated by carbon sources. Biotechnol Lett 42: 613–624.

27. Numazawa R, Tanaka Y, Nishioka S, Tsuji R, Maeda H, Tanaka M, Takeuchi M, Yamagata Y. 2024. *Aspergillus oryzae* PrtR alters transcription of individual peptidase genes in response to the growth environment. Appl Microbiol Biotechnol 108: 90.

28. Kerkaert JD, Huberman LB. 2023. Regulation of nutrient utilization in filamentous fungi. Appl Microbiol Biotechnol 107: 5873–5898.

29. Tanaka M. 2024. Transcriptional and post-transcriptional regulation of genes encoding secretory proteins in *Aspergillus oryzae*. Biosci Biotechnol Biochem 88:381–388.

30. Tanaka M, Ito K, Matsuura T, Kawarasaki Y, Gomi K. 2021. Identification and distinct regulation of three di/tripeptide transporters in *Aspergillus oryzae*. Biosci Biotechnol Biochem 85: 452–463.

31. Dantuma NP, Lindsten K, Glas R, Jellne M, Masucci MG. 2000. Short-lived green fluorescent proteins for quantifying ubiquitin/proteasome-dependent proteolysis in living cells. Nat Biotechnol 18: 538–543.

32. Heessen S, Dantuma NP, Tessarz P, Jellne M, Masucci MG. 2003. Inhibition of ubiquitin/proteasome-dependent proteolysis in *Saccharomyces cerevisiae* by a Gly-Ala repeat. FEBS Lett 555: 397–404.

33. Kobayashi T, Maeda H, Takeuchi M, Yamagata Y. 2017. Deletion of *admB* gene encoding a fungal ADAM affects cell wall construction in *Aspergillus oryzae*. Biosci Biotechnol Biochem 81: 1041–1050.

34. Oda K, Kakizono D, Yamada O, Iefuji H, Akita O, Iwashita K. 2006. Proteomic analysis of extracellular proteins from *Aspergillus oryzae* grown under submerged and solid-state culture conditions. Appl Environ Microbiol 72: 3448–3457.

35. Murakami K, Ishida Y, Masaki A, Tatsumi H, Murakami S, Nakano E, Motai H, Kawabe H, Arimura H. Isolation and characterization of the alkaline protease gene of *Aspergillus oryzae*. Agric Biol Chem 55: 2807–2811.

36. Cheevadhanarak S, Renno DV, Saunders G, Holt G. 1991. Cloning and selective overexpression of an alkaline protease-encoding gene from *Aspergillus oryzae*. Gene 108: 151–155.

37. Tilburn J, Sarkar S, Widdick DA, Espeso EA, Orejas M, Mungroo J, Peñalva MA, Arst HN Jr. 1995. The *Aspergillus* PacC zinc finger transcription factor mediates regulation of both acid- and alkaline-expressed genes by ambient pH. EMBO J. 14: 779–90.

38. Mellado L, Calcagno-Pizarelli AM, Lockington RA, Cortese MS, Kelly JM, Arst HN Jr, Espeso EA. 2015. A second component of the SltA-dependent cation tolerance pathway in *Aspergillus nidulans*. Fungal Genet Biol 82: 116–128.

39. Picazo I, Espeso EA. 2024. Interconnections between the Cation/Alkaline pH-Responsive Slt and the Ambient pH Response of PacC/Pal Pathways in *Aspergillus nidulans*. Cells. 13: 651.

40. Jin FJ, Watanabe T, Juvvadi PR, Maruyama J, Arioka M, Kitamoto K. 2007. Double disruption of the proteinase genes, tppA and pepE, increases the production level of human lysozyme by *Aspergillus oryzae*. Appl Microbiol Biotechnol 76: 1059–1068.

41. Zhu L, Nemoto T, Yoon J, Maruyama J, Kitamoto K. 2012. Improved heterologous protein production by a tripeptidyl peptidase gene (*AosedD*) disruptant of the filamentous fungus *Aspergillus oryzae*. J Gen Appl Microbiol 58: 199–209.

42. Maeda H, Sakai D, Kobayashi T, Morita H, Okamoto A, Takeuchi M, Kusumoto K, Amano H, Ishida H, Yamagata Y. 2016. Three extracellular dipeptidyl peptidases found in *Aspergillus oryzae* show varying substrate specificities. Appl Microbiol Biotechnol 100: 4947–4958.

43. Ridenour JB, Smith JE, Hirsch RL, Horevaj P, Kim H, Sharma S, Bluhm BH. 2014. *UBL1* of *Fusarium verticillioides* links the N-end rule pathway to extracellular sensing and plant pathogenesis. Environ Microbiol. 16: 2004–2022.

44. Maitland MER, Lajoie GA, Shaw GS, Schild-Poulter C. 2022. Structural and Functional Insights into GID/CTLH E3 Ligase Complexes. Int J Mol Sci. 23: 5863.

45. Lamb HK, Leslie K, Dodds AL, Nutley M, Cooper A, Johnson C, Thompson P, Stammers DK, Hawkins AR. 2003. The negative transcriptional regulator NmrA discriminates between oxidized and reduced dinucleotides. J Biol Chem 278: 32107–32114.

46. Wong KH, Hynes MJ, Todd RB, Davis MA. 2007. Transcriptional control of *nmrA* by the bZIP transcription factor MeaB reveals a new level of nitrogen regulation in *Aspergillus nidulans*. Mol Microbiol 66: 534–51.

47. Zhao X, Hume SL, Johnson C, Thompson P, Huang J, Gray J, Lamb HK, Hawkins AR. 2010. The transcription repressor NmrA is subject to proteolysis by three *Aspergillus nidulans* proteases. Protein Sci 19:1405–14019.

48. Li A, Parsania C, Tan K, Todd RB, Wong KH. 2021. Co-option of an extracellular protease for transcriptional control of nutrient degradation in the fungus *Aspergillus nidulans*. Commun Biol 4:1409.

49. Orejas M, Espeso EA, Tilburn J, Sarkar S, Arst HN Jr, Peñalva MA. 1995. Activation of the *Aspergillus* PacC transcription factor in response to alkaline ambient pH requires proteolysis of the carboxy-terminal moiety. Genes Dev 9: 1622–1632.

50. Peñalva MA, Arst HN Jr. 2002. Regulation of gene expression by ambient pH in filamentous fungi and yeasts. Microbiol Mol Biol Rev 66: 426–446.

51. Arst HN, Peñalva MA. 2003. pH regulation in *Aspergillus* and parallels with higher eukaryotic regulatory systems. Trends Genet 19: 224–231.

52. Peñalva MA, Tilburn J, Bignell E, Arst HN Jr. 2008. Ambient pH gene regulation in fungi: making connections. Trends Microbiol 16: 291–300.

53. Peñalva MA, Lucena-Agell D, Arst HN Jr. 2014. Liaison alcaline: Pals entice non-endosomal ESCRTs to the plasma membrane for pH signaling. Curr Opin Microbiol 22: 49–59.

54. Chen L, Zou G, Zhang L, de Vries RP, Yan X, Zhang J, Liu R, Wang C, Qu Y, Zhou Z. 2014. The distinctive regulatory roles of PrtT in the cell metabolism of *Penicillium oxalicum*. Fungal Genet Biol 63: 42–54.

55. Mizutani O, Kudo Y, Saito A, Matsuura T, Inoue H, Abe K, Gomi K. 2008. A defect of LigD (human Lig4 homolog) for nonhomologous end joining significantly improves efficiency of gene-targeting in *Aspergillus oryzae*. Fungal Genet Biol 45:878–889.

56. Suzuki K, Tanaka M, Konno Y, Ichikawa T, Ichinose S, Hasegawa-Shiro S, Shintani T, Gomi K. 2015. Distinct mechanism of activation of two transcription factors, AmyR and MalR, involved in amylolytic enzyme production in *Aspergillus oryzae*. Appl Microbiol Biotechnol 99:1805–1815.

57. Motohashi K. 2015. A simple and efficient seamless DNA cloning method using SLiCE from *Escherichia coli* laboratory strains and its application to SLiP site-directed mutagenesis. BMC Biotechnol 15:47.

58. Ichinose S, Tanaka M, Shintani T, Gomi K. 2014. Improved α-amylase production by *Aspergillus oryzae* after a double deletion of genes involved in carbon catabolite repression. Appl Microbiol Biotechnol 98:335–343.

59. Gomi K, Iimura Y, Hara S. 1987. Integrative Transformation of *Aspergillus oryzae* with a Plasmid Containing the *Aspergillus nidulans argB* Gene. Agric Biol Chem 51:2549–2555.

60. Maruyama J, Kitamoto K. 2008. Multiple gene disruptions by marker recycling with highly efficient gene-targeting background (Delta*ligD*) in *Aspergillus oryzae*. Biotechnol Lett 30, 1811–1817.

61. Laemmli UK. 1970. Cleavage of Structural Proteins during the Assembly of the Head of Bacteriophage T4. 5259. Nature 227:680–685.

